# A new local covariance matrix estimation for the classification of gene expression profiles in RNA-Seq data

**DOI:** 10.1101/766402

**Authors:** Necla Koçhan, Gözde Yazgı Tütüncü, Göknur Giner

## Abstract

**Background and Objective:** Recent developments in the next-generation sequencing (NGS) based on RNA-sequencing (RNA-Seq) allow researchers to measure the expression levels of thousands of genes for multiple samples simultaneously. In order to analyze these kind of data sets, many classification models have been proposed in the literature. Most of the existing classifiers assume that genes are independent; however, this is not a realistic approach for real RNA-Seq classification problems. For this reason, some other classification methods, which incorporates the dependence structure between genes into a model, are proposed. qtQDA proposed by Koçhan et al. [1] is one of those classifiers, which estimates covariance matrix by Maximum Likelihood Estimator.

**Methods:** In this study, we use a another approach based on local dependence function to estimate the covariance matrix to be used in the qtQDA classification model. We investigate the impact of different covariance estimates on RNA-Seq data classification.

**Results:** The performances of qtQDA classifier based on two different covariance matrix estimates are compared over two real RNA-Seq data sets, in terms of classification error rates. The results show that using local dependence function approach yields a better estimate of covariance matrix and increases the performance of qtQDA classifier.

**Conclusion:** Incorporating the true/accurate covariance matrix into the classification model is an important and crucial step particularly for cancer prediction. The local covariance matrix estimate allows researchers to classify cancer patients based on gene expression profiles more accurately. R code for local dependence function is available at https://github.com/Necla/LocalDependence.

## 1. Introduction

Dependence relation between random variables is one of the most commonly studied subjects in statistical data analysis. It is an important task to figure out the dependence structure of a data set and incorporate it into a statistical model in data analysis field. Generally, one can incorporate the dependence structure via covariance matrices which play an important role in multivariate statistical models, data classification, image processing, etc. A simple way to estimate the covariance matrix is to use Maximum Likelihood Estimator. However, this simple estimator may not reflect the complex dependence structures in medical and biological sciences due to the high dependence between the variables (attributes) in data sets. Hence, there have been a few recent approaches proposed for improving the covariance matrix estimation [2, 3] in the literature.

Caefer and Rotman [4] developed a quasi-local covariance matrix estimation to be applied on spectral data analysis. Instead of estimating the whole covariance matrix they use the variance of neighbours surrounding the reference point and they define dependence areas. That is, the points in highly variable areas will have higher variances and the points in low variable areas will have less variances, accordingly. Similar to Caefer and Rotman’s approach [4], Oruc and Ucer [5] proposed a new methodology to construct local dependence map which can identify three regions: positive, negative and zero dependence. They applied it on real medical data sets and showed that local dependence is much more informative in some instances.

Since it is known that RNA-Seq data sets are composed of many genes which are highly correlated with a high dependence degree, we claim that new samples will have an individual impact on the estimation of the covariance matrix while classifying the new samples. For this purpose, in this study, we propose a new type of covariance matrix estimate, which is called local covariance matrix, that can be implemented in qtQDA classifier. Integrating this new local covariance matrix into the qtQDA classifier improves the performance of the classifier. In this study, since the local covariance is updated for each new sample observation with a newly proposed method, the classifier, qtQDA, becomes an adaptive algorithm and we call it Local-quantile transformed Quadratic Discriminant Analysis (L-qtQDA).

## 2. Methodology

Classification of RNA-Seq data has become an important research area in the last decade. Particularly in cancer research, true classification of the sub-type of a patient with a particular cancer, leads a better predictive and a customized treatment for that patient. Therefore, classification of a patient to a cancer sub-type at gene expression level has a crucial importance. Due to the discrete structure of RNA-Seq data, classification of these kind of data is not as simple as other classification models that are proposed for continuous data types.

There are certain number of classifiers proposed especially for RNA-Seq data in the literature [6]. The most recent one is qtQDA classifier proposed by Koçhan et al. [1]. Since qtQDA incorporates the dependence structure into the model, we apply qtQDA in order to compare a differently estimated covariance matrix, local covariance matrix, with the simple one used in qtQDA model. In the following section we explain the qtQDA classifier in details.

### 2.1. Negative Binomial Marginals

Suppose that we have *k* distinct classes and want to classify new samples into one of those *k* classes on the basis of *m* genes. Let 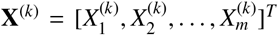 be a gene expression data matrix from *k*th class where 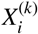 is the number of reads (counts) for gene *i*. Assume that counts are marginally negative binomial distributed, i.e.

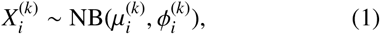

where 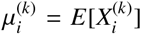 and 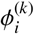 is the dispersion for gene *i*. It can be easily calculated that

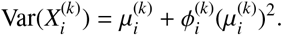

If 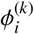 is different than zero then

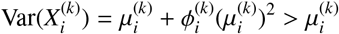

which is consistent with known properties of RNA-seq data when there are biological replicates (readers are referred to [7] for more details).

### 2.2. Quantile Transformation

In order to incorporate the dependence into the model, a quantile transformation process is applied:

1. Let **Z**^(*k*)^ be an *m*-vector from a multivariate normal distribution: **Z**^(*k*)^ ~ MVN(**0**, Σ^(*k*)^), where 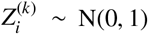.
2. Then transform *i*th component of **Z**^(*k*)^ into the *i*th component of **X**^(*k*)^

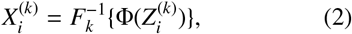

where Φ is the standard normal distribution function, **X**^(*k*)^ is the transformed random variable and *F*_*k*_ is the 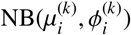 distribution function.

Note here that each class has its own different covariance matrix which is expected to increase the performance of the classification.

### 2.3. Classification

Suppose we observe a new sample 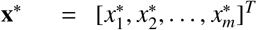 from unknown class *y*\*, where *y*\* ∈ {1, 2,…, *K*} is the class label. Using inverse of quantile transformation, we transform components of the new sample **x**\* to a new vector **z**\*^(*k*)^ which is multivariate normally distributed with parameters µ = **0** and Σ = Σ^(*k*)^. It is obvious that this transformation is applied for each class separately. Then, by Bayes theorem, posterior probability of **x**\* belonging to the *k*th class is given as

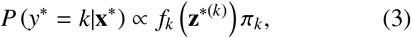

where π_*k*_ is the prior probability and *f*_*k*_ is the density

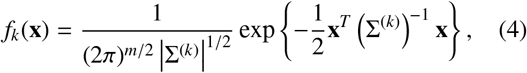

Using Eq. (3) and Eq. (4), the quadratic discriminant score for qtQDA can be defined as follows:

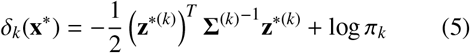

Thus we classify new sample **x**\* into one of *k* distinct classes which maximizes the Eq. (5).

### 2.4. Parameter Estimation

In order to apply the model in practice, there exist some parameters to be estimated in the classification model. Now, we explain how they are estimated.

- *Negative Binomial Parameters(mean and dispersion)*. Like qtQDA, we use **estimateDisp** function in the R package **edgeR**. This function estimates mean using maximum likelihood and calculates a matrix of likelihoods for each gene at a set of dispersion grid points. Then weighted likelihood empirical Bayes method is applied to obtain posterior dispersion estimates for each gene [7, 8].
- *Covariance Matrix*. We apply quantile transformation to produce multivariate normally distributed vectors to be used in evaluating the posterior probabilities. To do so, we need to estimate the covariance matrix for each class. Unlike Koçhan et al. [1], in this study we use local dependence function explained in Section 3 in order to improve the covariance matrix estimation and we call this estimation as local covariance matrix. Note here that similar to Koçhan et al., we use the R package “corpcor” to guarantee that local covariance matrix is symmetric and positive definite for downstream analysis.

## 3. Local Dependence Function

Let (*X*, *Y*) be a continuous bivariate random variable with joint cumulative distribution function *F* (*x*, *y*) and with joint probability density function *f* (*x*, *y*). Then the Pearson correlation coefficient between *X*, *Y* is given as

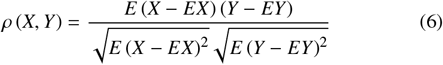

Indeed, Eq. (6) is a way of measuring linear dependence between two random variables and in some researches it is called measure of association [9, 10]. But in some cases, this strength of association between two random variables can vary locally. In order to define a local measure of the association between two random variables Bairamov and Kotz [9] proposed a new local dependency function which replaces the expectations *EX* and *EY* by conditional expectations *E* (*X*|*Y = y*) and *E* (*Y*|*X* = *x*), respectively. The Bairamov & Kotz (2000) local dependence function is given as follows:

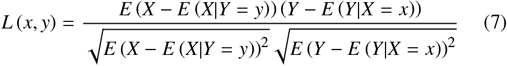

Let ε_*X*_ (*y*) = *EX* − *E* (*X*|*Y* = *y*) and ε_*Y*_ (*x*) = *EY* − *E* (*Y*|*X* = *x*). Then

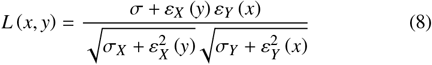

where σ = *Cov* (*X*, *Y*).

Thus, local dependence function *L* (*x*, *y*) which represents the dependence between *X* and *Y* at any specific point (*x*, *y*) is more robust and accurate if there exists a dependence in the model.

In order to estimate the covariance matrix from the data available we need to estimate the local dependence function from the data. Therefore, Nadaraya [11] and Watson [12] proposed the following estimates for the regression functions *E* (*X*|*Y* = *y*) and *E* (*Y*|*X* = *x*):

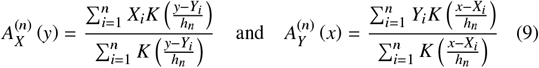

where *K* is an integrable kernel function with short tails and *h*_*n*_ → 0 is a width sequence tending zero at approximate rates.

Since it is given in [13] that the optimal choice for *h* is

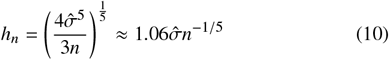

where 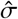 is the standard deviation of the samples, we use Eq. (10) in order to estimate the conditional expectations. Moreover, we use triangular kernel function which is given as follows:

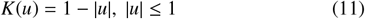

Using those estimates given in Eq. (9), we suggest the following estimate for local dependence function

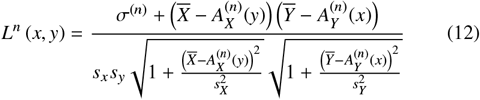

where

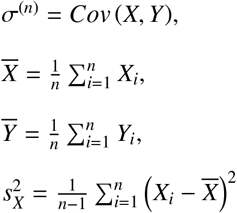

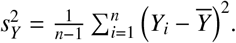

Note here that *X* and *Y* are any two genes across samples.

## 4. Results

In this section, we implement qtQDA based on two different estimates of covariance matrix. For qtQDA classifier, we use R package “qtQDA” which is available at https://github.com/goknurginer/qtQDA. For the discriminant function in the qtQDA package, We use trended dispersion estimate. Then, we compare the classification error rates using two real RNA-Seq data sets. These are not only publicly available data sets but also commonly used data sets in order to test the performance of RNA-Seq classification methods.

The first data is cervical cancer data (see [14]). The cervical cancer data is composed of 714 microRNAs and 58 samples where 29 samples are tumor and 29 samples are non-tumor.

The second data is HapMap data (see [15, 16]). Similar to cervical cancer data, the HapMap data also includes two groups of samples; CEU and YRI where CEU represents Utah residents with Northern and Western European Ancestry and YRI represents Yoruba in Ibadan and Nigeria, respectively. There are 91 CEU samples and 89 YRI samples with a total number of 52,580 genes.

It is known that RNA-Seq technology measures the expression levels of thousands of genes for multiple samples. However, not all genes are relevant and informative. Therefore, a gene selection technique is required not only to reduce the computing time but also to improve the classification performance. We apply edgeR pipeline to select informative genes which will be used in the classification algorithm. Basically, a likelihood ratio test (LRT) is performed in edgeR to detect differentially expressed (DE) genes between groups. After that, DE genes are sorted according to the value of LRT statistic and finally, the top *m* genes are used for the classification process. In our study, the top 20, 50, 100, 200, 300, 500 DE genes are selected for both cervical cancer data and HapMap data.

After conducting gene selection procedure, we randomly split the dataset into two sets: training set and test set. 70% of the dataset is randomly assigned to the training and the rest 30% of the dataset is assigned to the test set. Training set is used to train the classifiers and test set is used to measure the misclassification rate. The whole procedure is repeated 300 times for different number of genes and the average misclassification rate is computed.

The comparison results are given Table 1. It is obvious to see that improving the covariance matrix estimate, i.e using local dependence function to estimate the covariance matrix, leads generally better results. Interestingly, for both data sets, qtQDA performs better then L-qtQDA. However, for the cervical cancer data, we obtain better performances except for 20 and 200 genes selected in gene selection process. For HapMap data, we obtain better performances except for 200 and 500 genes selected in gene selection process. Overall we can conclude that L-qtQDA performs generally better than qtQDA.

**Table 1.** Classification error rates for cervical cancer and HapMap data sets

| Data | # of genes | qtQDA | L-qtQDA |
| --- | --- | --- | --- |
| Cervical | 20 | **0.0367** | 0.0372 |
|  | 50 | 0.0280 | **0.0265** |
|  | 100 | 0.0126 | **0.0124** |
|  | 200 | **0.0117** | 0.0122 |
|  | 300 | 0.0161 | **0.0159** |
|  | 500 | 0.0189 | **0.0170** |
| HapMap | 20 | 0.0172 | **0.0166** |
|  | 50 | 0.0064 | **0.0057** |
|  | 100 | 0.0448 | **0.0434** |
|  | 200 | **0.0120** | 0.0116 |
|  | 300 | 0.0074 | **0.0073** |
|  | 500 | **0.0106** | 0.0109 |

## 5. Discussion

It is shown in qtQDA paper that instead of assuming independency of genes, incorporating the dependence between genes into the model can lead better performance [1]. Thus, in this paper, we investigated a new estimate for covariance matrix that can capture a true dependence structure and then can be implemented in qtQDA. We anticipated the new covariance matrix as local dependency of genes as the degree of dependency between pairs of genes may vary and affect the estimation of covariance matrix. We showed that estimating covariance matrix locally can lead even better performance in the RNA-Seq data classification. Therefore, one should take the estimation of the covariance matrix into account since it has a significant effect on the classification performances.

Since we only used triangular kernel function and Gaussian banwidth in local dependency calculation, we note here that different kernel functions and an optimal banwidth selection can also be implemented and may improve the classification performances. The only disadvantage of the L-qtQDA is that the algorithm is computationally intensive due to the estimation of the local covariance matrix.

## 6. Conclusion

In this study we investigated the impact of covariance matrix estimated with the help of local dependence function on RNA-Seq data classification. This new approach assumes the dependencies between genes are locally defined rather than complete dependency. We have shown that locally estimated covariance matrix is more effective than simple covariance matrix on real RNA-Seq data classification. We believe that this new estimation technique will be useful for classification of RNA-Seq profiles or other genomic studies.

